# Characterizing an electronic–robotic targeting platform for precise and fast brain stimulation with multi-locus transcranial magnetic stimulation

**DOI:** 10.1101/2024.03.12.584601

**Authors:** Renan H. Matsuda, Victor H. Souza, Thais C. Marchetti, Ana M. Soto, Olli-Pekka Kahilakoski, Mikael Laine, Heikki Sinisalo, Dubravko Kicic, Pantelis Lioumis, Risto J. Ilmoniemi, Oswaldo Baffa

## Abstract

1.

**Background:** Multi-locus TMS (mTMS) enables precise electronic control of brain stimulation targeting, eliminating the need for physical coil movement. However, with a small number of coils, the stimulation area is constrained, and manually handling the coil array is cumbersome. Combining electronic mTMS targeting with robotics will enable automated, user-independent, and precise brain stimulation protocols.

**Objective:** Characterizing an open-source electronic–robotic mTMS platform for rapid and accurate brain stimulation targeting.

**Methods:** We developed an automated robotic mTMS positioning platform. The accuracy of the system was quantified with a TMS characterizer that measures the TMS-induced electric field on a spherical cortex model. We used a 5-coil mTMS device equipped with a set of five coils coupled to a collaborative robot. The induced electric-field distortion generated by robot coupling was evaluated for each coil. We compared the accuracy of robotic–electronic targeting by repositioning the mTMS coil set with the robotic and the conventional manual positioning.

**Results:** Our collaborative robot-based system offers submillimeter precision and autonomy in positioning mTMS coil sets. The electronic–robotic mTMS platform was approximately 1.8 mm and 1.0° more accurate than the conventional manual positioning. Integrating robotics and mTMS automates brain stimulation procedures, resulting in minimal reliance on user expertise and subjective analysis.

**Conclusion:** Our open-source platform combining rapid mTMS targeting with robotic precision enhances the safety and reproducibility of brain stimulation techniques, enabling more efficient and reliable outcomes than previous techniques.

## 3. Introduction

In order to enhance the safety and effectiveness of non-invasive brain stimulation, it is imperative to adopt automation and precise targeting of cortical structures. Multi-locus transcranial magnetic stimulation (mTMS) presents a significant advance, allowing electronic stimulation of nearby cortical regions without the need for physical movement of the stimulation coil (Nieminen *et al*., 2022; Souza *et al*., 2022). This technology allows interactions with local cortical networks at millisecond and millimeter scales (Nieminen *et al*., 2019; Tervo *et al*., 2020; Sinisalo *et al*., 2023). Despite the promising aspects of multi-locus TMS, existing coil sets encounter two primary challenges. First, mTMS has a limited electronic targeting range, in our case with 5 coils, a 30-mm diameter region. Second, the coil array assembly can be heavy, weighing over 5 kg for a 5-coil set with cables (Sinisalo *et al*., 2023). Consequently, the manual placement of the coil set on the scalp is a slow and physically demanding process. These limitations restrict the application of automated algorithms, confining them to single predetermined areas and demanding highly trained and physically capable personnel for manipulating the coil sets. Collaborative robots have demonstrated the ability to ease the process of TMS coil placement, improving the reproducibility and accuracy of reaching the desired cortical target (Goetz *et al*., 2019). These robots can autonomously compensate for patients’ head movements. However, existing robotized TMS applications do not seamlessly integrate with mTMS algorithms and have limited velocities (around 20 cm/s) to safely adjust the stimulation target for human applications (Kantelhardt *et al*., 2010). Additionally, existing commercial robotic TMS solutions rely on closed-source platforms associated with specific robotic arms, leading to potential cost constraints and implementation difficulties. These factors prevent researchers from incorporating novel algorithms as needed. Recently, we developed an open-source electronic–robotic control that seamlessly integrates rapid mTMS electronic targeting with precise and autonomous robotic handling. Our system eliminates the need for manual manipulation of mTMS coil sets, significantly improving the safety and precision of the technique. The platform is readily accessible at https://github.com/biomaglab/tms-robot-control.

Our goal was to characterize the accuracy and precision in targeting the induced electric field (E-field) by combining the mTMS electronic control and the automated robotic positioning. We also evaluated the distortion on the E-field caused by attaching mTMS coil sets to the robot flange and compared the manual and robotic– electronic brain targeting. This platform opens up exciting possibilities for developing new brain stimulation paradigms, such as closed-loop and operator-independent protocols capable of effectively covering large cortical brain areas.

## 4. Material and Methods

We characterized our platform utilizing the Elfin E5 collaborative robot (Han’s Robot Co Ltd, China). This robot has 6 joints, a 5-kg payload capacity, an operational range of up to 80 cm, and a repeatability accuracy of ± 0.05 mm. The electronic–robotic control was developed to operate with the open-source neuronavigation system InVesalius Navigator (Souza *et al*., 2018). We used an mTMS device with a 5-coil set and an effective stimulation range of 30 mm in diameter. The 5-coil set comprises two four-leaf-clover coils, two figure-of-eight coils, and an oval coil (Nieminen *et al*., 2022). We utilized our TMS characterizer to measure the induced E-field spatial distribution on a 70-mm-radius spherical cortical model (Nieminen, Koponen and Ilmoniemi, 2015). Bottom of the mTMS coil set was placed 15 mm above the measurement probe.

### Electronic–robotic targeting

The integration between the robotic control and the mTMS electronic targeting was implemented in InVesalius. We developed a user interface to define the desired scalp targets for the mTMS coil set position and orientation. Given these coil-set locations on the scalp, the user defines the cortical targets for the mTMS within a 30-mm diameter region around the center of the mTMS coil set. The cortical targets are linearly projected at a 15-mm depth (Koponen, Nieminen and Ilmoniemi, 2018) along the coil’s normal axis from the scalp target following the spherical brain model used to design the 5-coil sets (Koponen, Nieminen and Ilmoniemi, 2015; Nieminen *et al*., 2022; Souza *et al*., 2022).

### Experimental characterization

We characterized the electronic–robotic targeting in three steps: 1) we verified the E-field distortion caused by attaching the mTMS coil sets to the robot flange. We measured the induced E-fields of each of the five mTMS coils individually, with and without the robot attached to the coil assembly. The measurement of the induced E-field had 250 data points in a 40-mm diameter circular area from the central point of the mTMS transducer on the spherical cortical model. We used a wooden stand below the mTMS coil set to ensure it had the same position relative to the TMS characterizer with and without the robot attached (figure 1). First, we measured with the mTMS coil set screwed to the robot flange. Then, the robotic arm was detached and moved at least 50 cm away from the measurement platform.

**Figure 1.**
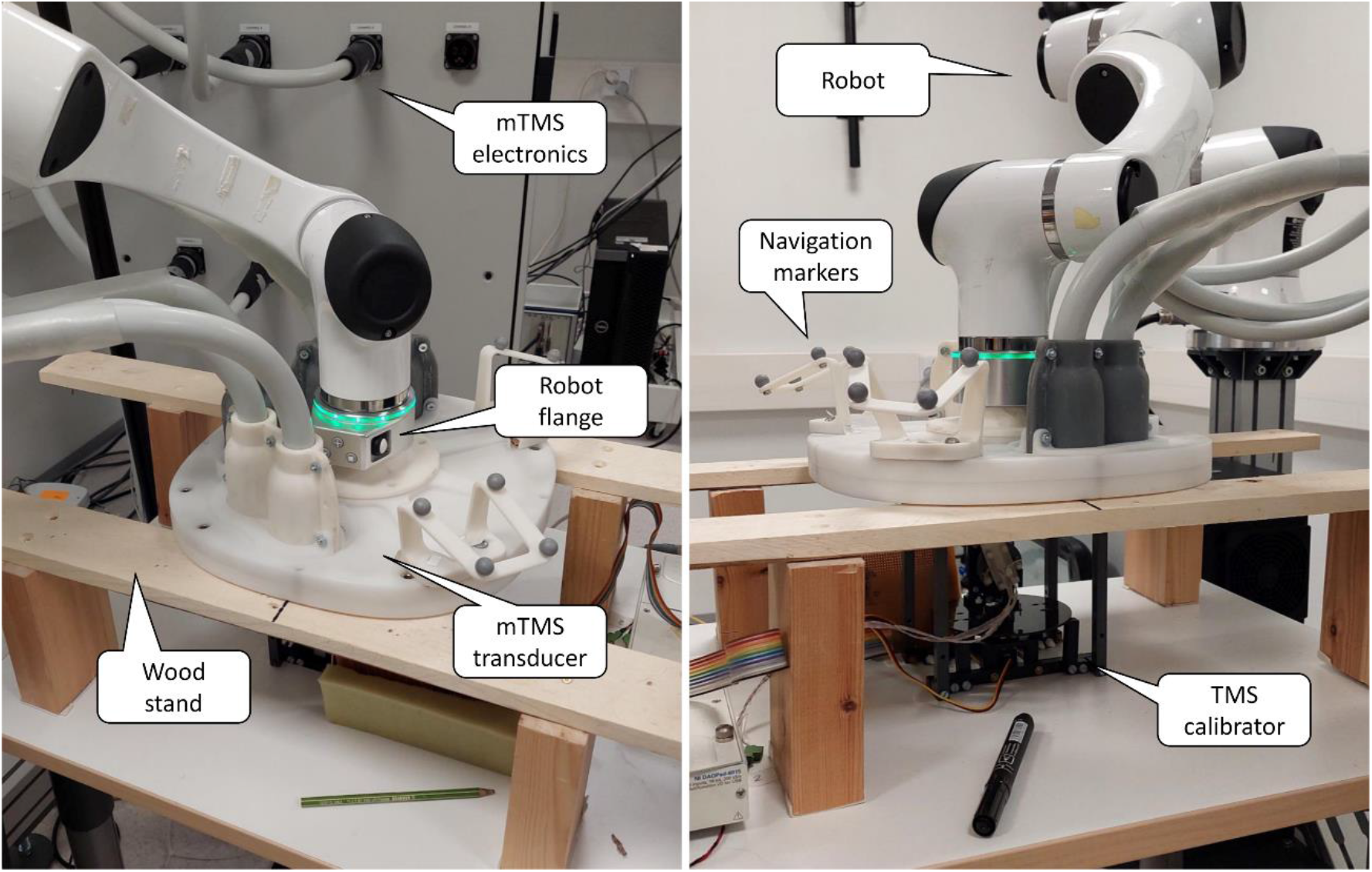
Two views from the experimental setup with the mTMS transducer held by the robot on top of the TMS calibrator

Next, 2) we characterized the robotic–electronic targeting accuracy by physically moving the coil set and compensating for this displacement with the electronic targeting algorithm. We created 10 mTMS scalp targets within a 30-mm diameter circle of the calibrator center point; the rotation angle was selected in the range from −30° to 30°. Scalp target locations and orientations were pseudo-randomly sampled from a uniform distribution. For tracking the coil set, we used a motion capture system with eight Flex13 cameras (OptiTrack, NaturalPoint, Inc., USA) together with the InVesalius neuronavigation system (Souza *et al*., 2018). Neuronavigation was performed with a magnetic resonance image (MRI) phantom with similar dimensions (11 × 15 × 17 cm3) to the TMS calibrator. The MRI phantom comprises a set of 180 images generated in MATLAB 2022a (The MathWorks Inc, USA). The image represents a conventional 3D T1-weighted structural MRI with 256 × 256 × 180 voxels of size 1 × 1 × 1 mm3, figure 2(a). The neuronavigation fiducials were defined at the bottom right and left and at the anterior top-middle of both phantom MRI and TMS calibrator, figure 2(b) and 2(c).

**Figure 2.**
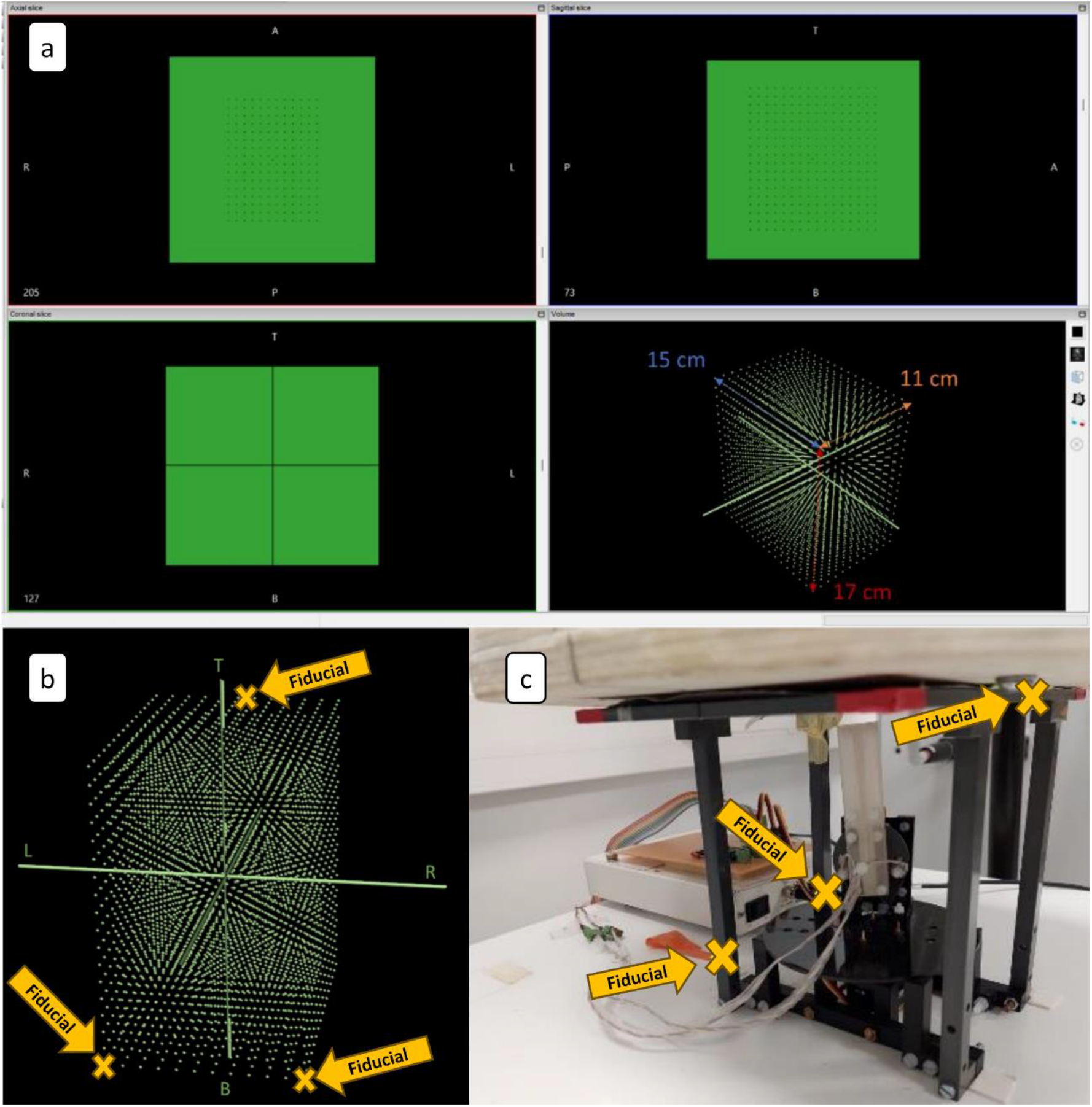
(a) InVesalius window with the MRI phantom with similar dimensions as the TMS calibrator. The green dots in the 3D panel are spaced by 1 cm in all axes. (b) The MRI phantom highlights the fiducials located at the bottom right and left and at the anterior top-middle of the MRI. (c) Photograph of the TMS calibrator with fiducials placed according to the MRI.

The robotized mTMS platform automatically estimates the offset to translate the maximum E-field from the center of the mTMS transducer to the center of the TMS calibrator for all scalp targets. The E-field for each target was measured at 100 points, covering a 30-mm diameter area centered on the TMS coil. The experiment was repeated three times, resulting in 30 pseudo-random scalp targets. The dispersion (standard deviation, *SD*) was computed from the *x* and *y* centroid coordinates of the focal E-field area (70% of the peak E-field (10)).

Last, 3) we evaluated the accuracy of repositioning the mTMS coil set over the scalp target in two mockup experiments: first, with the mTMS transducer positioned by the robot and second, manually. We defined a scalp target approximately above the motor cortex and in an initial arbitrary location far from a dummy head. We alternated the robot position 15 times between the target and the initial position. We used InVesalius connected to the tracking device Polaris Vega VT (Northern Digital Inc., Canada), to record the coil coordinates. Neuronavigation was performed with the Montreal Neurological Institute average MRI (Evans *et al*., 1993) based on 152 MRIs. To assess the difference in robotic and manual repositioning accuracy, we applied a two-way ANOVA followed by Tukey HSD post-hoc multiple comparisons. Statistical analysis was performed with custom scripts written in R 4.2 (R Core Team, Austria). The significance threshold was set at 0.05.

## 5. Results

### E-field distortion

The robot attached to the mTMS coil set did not cause any observable differences in the E-field spatial distribution of all coils. The maximum E-field recorded with the cobot attached had an average marginal increase of 3.3% ± 3.2% compared to without the cobot. Figure 3 depicts the E-field spatial distribution from the five mTMS coils.

**Figure 3.**
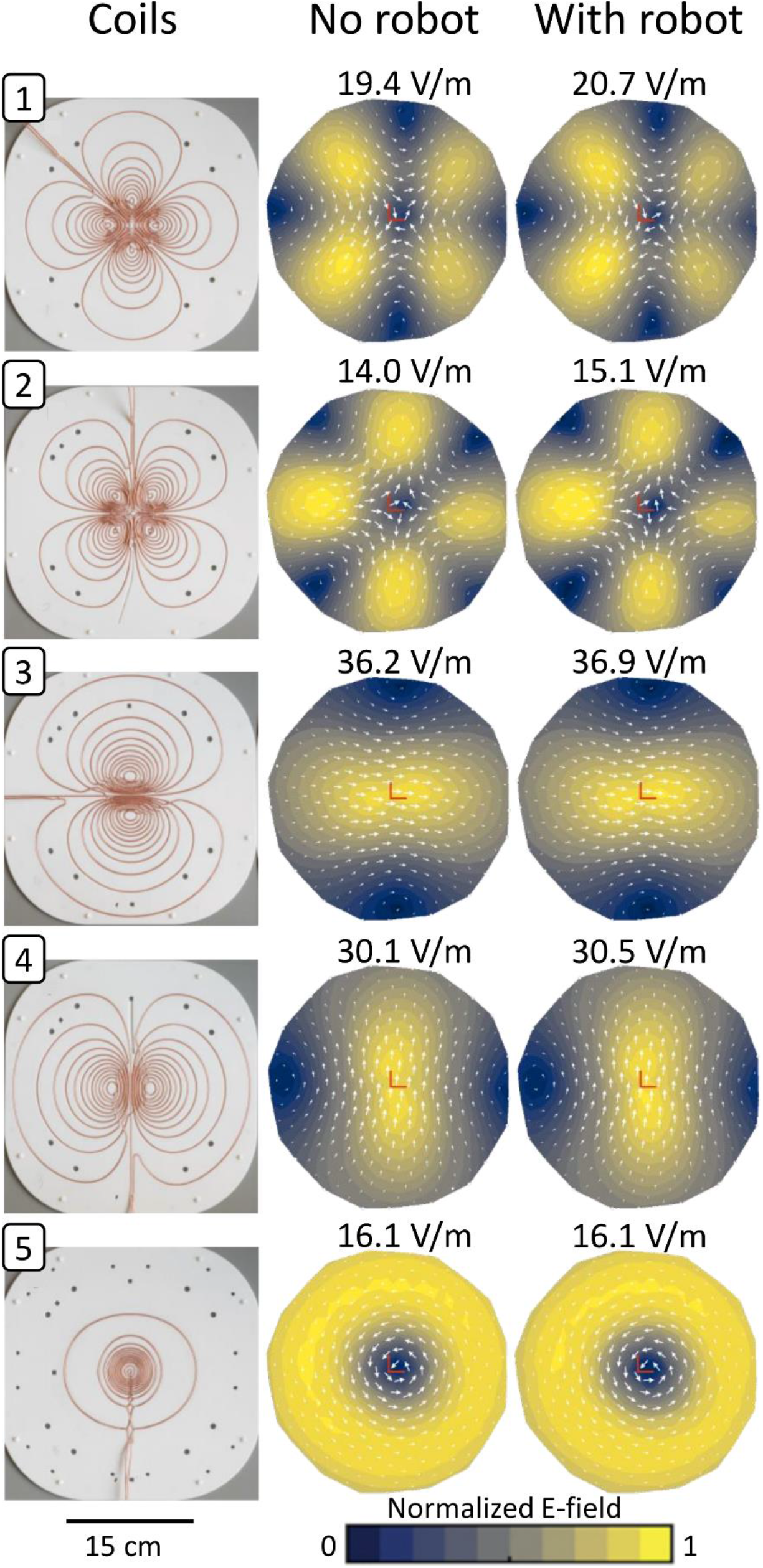
The coil windings and their corresponding E-field distribution with and without the robot. The values on each E-field distribution represent the maximum E-field intensity.

### Robotic–electronic cortical targeting accuracy

The induced E-field centroid dispersion of the autonomous robotized mTMS in the *x* and *y* axes were ± 0.7 and ± 0.2 mm, respectively. The dispersion of the peak E-field intensity was ± 1.0 V/m. The average E-field distribution across all measurements is illustrated in figure 4(a).

**Figure 4.**
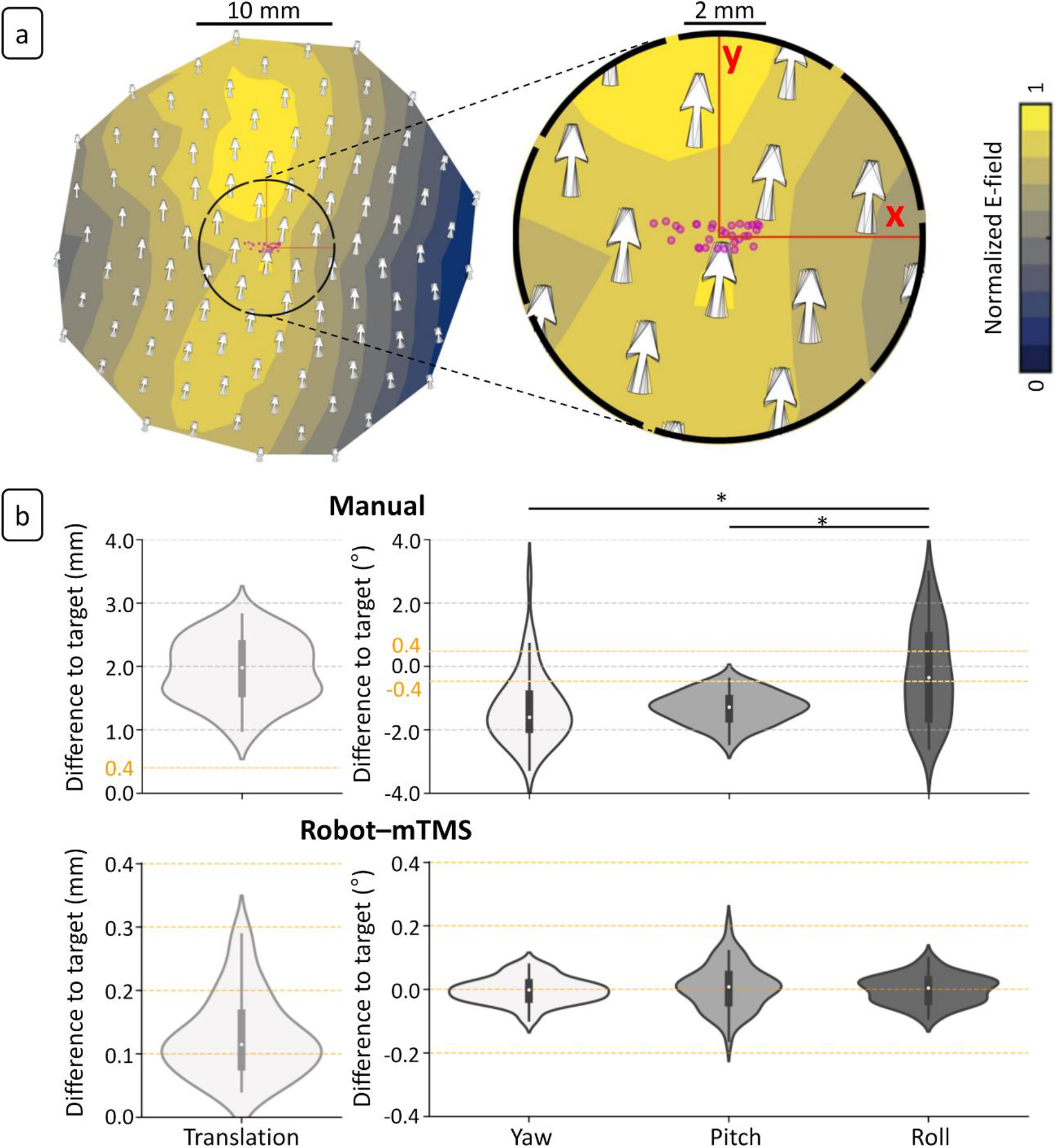
Experimental characterization and demonstration of robotic–electronic mTMS. (a) E-field dispersion of the robotized mTMS. The purple circles are the centroids of the E-field distributions for each mTMS target. The red axes indicate the center of the TMS calibrator, and the white arrows show the E-field orientation at a given calibrator’s probe position. The arrows are overlapped over each mTMS target. The black dashed circle (10-mm diameter) indicates the spatial extent of the zoomed panel on the right side. (b) Measured accuracy in repositioning the mTMS coil set on the target manually (top panel) and with the robotic control (bottom panel). ^*^ *p* < 0.001.

### Comparison of manual and robotic–electronic cortical target positioning

The average robotized positioning was about 1.8 mm and 1.0° more accurate than the manual positioning for the translation (F_1,95_ = 646.6; *p* < 0.001) and rotation angles (F_1,289_ = 87.57; *p* < 0.001), respectively. The average distance to the target with the robotized repositioning was 0.3 mm and with orientations within −0.2° to 0.2°. No difference was found between the rotation angles (F_2,141_ = 646.6; *p* = 0.638). The accuracy for manual repositioning was about 3.0 mm and ± 3.0°. There was a significant difference within the rotation angles (F_2,144_ = 16.52; *p* < 0.001). The roll was 1.1° higher than the pitch (*p* < 0.001) and 1.1° smaller than the yaw (*p* < 0.001) (figure 4(b)).

## 6. Discussion

We harnessed the high accuracy and autonomous capabilities of a collaborative robot to enable hands-free and precise positioning of mTMS coil sets. We demonstrated that attaching the mTMS coil set to the robot flange does not alter the E-field distribution because, if the robot coupling had affected the E-field due to mutual inductance, the E-field would have been decreased. Interestingly, the maximum E-fields are slightly higher or equal with the robot coupling than without the coupling. These increases may be caused by the robot weight applied in the wood stand that can press the coil set slightly closer to the TMS calibrator; even a small variation will cause an observable increase in the maximum resulting E-fields. Our findings indicate that the robotized–electronic system achieves superior accuracy compared to manual positioning and demonstrates comparable stability and accuracy to existing robotized TMS systems (Richter *et al*., 2010; Pennimpede *et al*., 2013; Grab *et al*., 2018; Goetz *et al*., 2019; Noccaro *et al*., 2021). Our open-source platform for robotic–electronic targeting can be used to automate TMS protocols, including stimulation target, hotspot identification, and precise motor mapping, employing closed-loop algorithms while minimizing reliance on user experience and subjective analysis (Harquel *et al*., 2017; Tervo *et al*., 2020; Weise *et al*., 2023).

A limitation of our robot control implementation is that the transformation between the robot to the tracking device is based on the physical location of the robot base and the tracking cameras. Thus, displacements of any of these two components will nullify the transformation matrix, compromising the robot control accuracy. A multi-camera motion capture system, such as the one used in this study, alleviates this limitation with a fixed installation. Compared to compact 2-camera models, this setup has a larger coverage volume that is less affected by the occlusion of a single camera’s field-of-view and might provide more stable tracking when moving the coil set to opposing sides of the scalp, for instance, from occipital to anterior lobes. Additionally, head marker displacements cause shifts in the placement of the robotic control, leading to inaccuracies in brain stimulation. To overcome this limitation in the future, markerless head pose estimation software can be utilized (Matsuda *et al*., 2023). This would add more freedom to the subjects and create a more natural experimental situation.

## 7. Conclusion

The development of our open-source platform, integrating electronic mTMS targeting and robotics significantly enhances the safety and reproducibility of brain stimulation procedures. We foresee that the implementation of automated and fast stimulation protocols fosters the creation of streamlined, user-independent, and highly efficient TMS procedures in both clinical and research settings. This capability will improve treatments and allow for more sophisticated stimulation protocols aimed at a better understanding of cerebral functions.

## 9. Acknowledgment

We are thankful to Lourenço Rocha, Carlos Renato Silva, and Fernando Henrique Torrieri for technical support.

## References

Evans, A. C. et al. (1993) ‘3D statistical neuroanatomical models from 305 MRI volumes’, 1993 IEEE Conference Record Nuclear Science Symposium and Medical Imaging Conference, pp. 1813–1817. doi: 10.1109/NSSMIC.1993.373602.

Goetz, S. M. et al. (2019) ‘Accuracy of robotic coil positioning during transcranial magnetic stimulation’, Journal of Neural Engineering, 16(5), p. 054003. doi: 10.1088/1741-2552/ab2953.

Grab, J. G. et al. (2018) ‘Robotic TMS mapping of motor cortex in the developing brain’, Journal of Neuroscience Methods, 309(August), pp. 41–54. doi: 10.1016/j.jneumeth.2018.08.007.

Harquel, S. et al. (2017) ‘Automatized set-up procedure for transcranial magnetic stimulation protocols’, NeuroImage, 153(April), pp. 307–318. doi: 10.1016/j.neuroimage.2017.04.001.

Kantelhardt, S. R. et al. (2010) ‘Robot-assisted image-guided transcranial magnetic stimulation for somatotopic mapping of the motor cortex: a clinical pilot study’, Acta Neurochirurgica, 152(2), pp. 333–343. doi: 10.1007/s00701-009-0565-1.

Koponen, L. M., Nieminen, J. O. and Ilmoniemi, R. J. (2015) ‘Minimum-energy coils for transcranial magnetic stimulation: Application to focal stimulation’, Brain Stimulation. Elsevier Inc., 8(1), pp. 124–134. doi: 10.1016/j.brs.2014.10.002.

Koponen, L. M., Nieminen, J. O. and Ilmoniemi, R. J. (2018) ‘Multi-locus transcranial magnetic stimulation—theory and implementation’, Brain Stimulation. Elsevier Ltd, 11(4), pp. 849–855. doi: 10.1016/j.brs.2018.03.014.

Matsuda, R. H. et al. (2023) ‘MarLe: Markerless estimation of head pose for navigated transcranial magnetic stimulation’, Physical and Engineering Sciences in Medicine, 46(2), pp. 887–896. doi: 10.1007/s13246-023-01263-2.

Nieminen, J. O. et al. (2019) ‘Short-interval intracortical inhibition in human primary motor cortex: A multi-locus transcranial magnetic stimulation study’, NeuroImage, 203(September). doi: 10.1016/j.neuroimage.2019.116194.

Nieminen, J. O. et al. (2022) ‘Multi-locus transcranial magnetic stimulation system for electronically targeted brain stimulation’, Brain Stimulation. The Author(s), 15(1), pp. 116–124. doi: 10.1016/j.brs.2021.11.014.

Nieminen, J. O., Koponen, L. M. and Ilmoniemi, R. J. (2015) ‘Experimental characterization of the electric field distribution induced by TMS devices’, Brain Stimulation. Elsevier Inc., 8(3), pp. 582–589. doi: 10.1016/j.brs.2015.01.004.

Noccaro, A. et al. (2021) ‘evelopment and validation of a novel calibration methodology and control approach for robot-aided Transcranial Magnetic Stimulation (TMS)’, 9294(c), pp. 1–12. doi: 10.1109/TBME.2021.3055434.

Pennimpede, G. et al. (2013) ‘Hot spot hound: A novel robot-assisted platform for enhancing TMS performance’, Proceedings of the Annual International Conference of the IEEE Engineering in Medicine and Biology Society, EMBS, pp. 6301–6304. doi: 10.1109/EMBC.2013.6610994.

Richter, L. et al. (2010) ‘Fast robotic compensation of spontaneous head motion during Transcranial Magnetic Stimulation (TMS)’, in UKACC International Conference on CONTROL 2010. Institution of Engineering and Technology, pp. 872–877. doi: 10.1049/ic.2010.0396.

Sinisalo, H. et al. (2023) ‘Modulating brain networks in space and time: Multi-locus transcranial magnetic stimulation’, Clinical Neurophysiology. doi: 10.1016/j.clinph.2023.12.007.

Souza, V. H. et al. (2018) ‘Development and characterization of the InVesalius Navigator software for navigated transcranial magnetic stimulation’, Journal of Neuroscience Methods, 309(June), pp. 109–120. doi: 10.1016/j.jneumeth.2018.08.023.

Souza, V. H. et al. (2022) ‘TMS with fast and accurate electronic control: Measuring the orientation sensitivity of corticomotor pathways’, Brain Stimulation. Elsevier Inc., 15(2), pp. 306–315. doi: 10.1016/j.brs.2022.01.009.

Tervo, A. E. et al. (2020) ‘Automated search of stimulation targets with closed-loop transcranial magnetic stimulation’, NeuroImage. Elsevier Inc., p. 117082. doi: 10.1016/j.neuroimage.2020.117082.

Weise, K. et al. (2023) ‘Precise motor mapping with transcranial magnetic stimulation’, Nature Protocols. Springer US, 18(2), pp. 293–318. doi: 10.1038/s41596-022-00776-6.

